# A transitory signaling center controls timing of primordial germ cell differentiation

**DOI:** 10.1101/2020.08.01.232298

**Authors:** Torsten U. Banisch, Maija Slaidina, Selena Gupta, Lilach Gilboa, Ruth Lehmann

## Abstract

Organogenesis requires exquisite spatio-temporal coordination of cell morphogenesis, migration, proliferation and differentiation of multiple cell types. For gonads, this involves complex interactions between somatic and germline tissues. During *Drosophila* ovary morphogenesis primordial germ cells (PGCs) are either sequestered in stem cell niches and maintained in an undifferentiated, germline stem cell state, or transition directly towards differentiation. Here, we identify a mechanism that links hormonal triggers of somatic tissue morphogenesis with PGC differentiation. An early ecdysone pulse initiates somatic swarm cell (SwC) migration, positioning them close to PGCs. A second hormone peak activates Torso-like signal in SwCs, which stimulates the Torso RTK signaling pathway in PGCs promoting their differentiation by de-repression of the differentiation gene *bag of* marbles. Thus, systemic temporal cues generate a transitory signaling center that coordinate ovarian morphogenesis with stem cell self-renewal and differentiation programs, a concept applicable broadly to the integration of stem cells and their niches.

**HIGHLIGHTS:**

- Steroid pulses coordinate gonadogenesis, stem cell self-renewal and differentiation
- An early steroid pulse initiates migration of Swarm Cells, a transitory support cell type
- A late steroid pulse induces Torso-like expression, a Torso receptor tyrosine kinase (RTK) activator, in Swarm Cells
- Torso RTK signaling in primordial germ cells activates the key differentiation gene *bam* by relieving Krüppel-mediated repression

## INTRODUCTION

Organ development and function requires complex interactions between cell types, which include the orchestration of proliferation rates, initiation of differentiation and tissue morphogenesis. Such widespread coordination typically requires systemic cues, e.g. hormones, that elicit specific cellular responses and wide-spread cell-cell signaling events at a given time point (Stamatiades and Kaiser, 2018; Yamanaka et al., 2013). During development, specialized groups of cells can aid in this coordinative effort by inducing cell fate decisions or patterning of surrounding cell types, and thus function as signaling or organizing centers (Anderson and Stern, 2016; Basson, 2012). While hormonal inputs and downstream signals are known to often act in a transient manner to allow for a stepwise progression of development, it is not well understood how cells perceive and integrate temporal cues and coordinate their response at an organ level.

Such higher-order regulation is exceedingly more complex if developing organs harbor stem cells or their precursors which need to be maintained in their naïve, undifferentiated state until their proper niches have formed and they transition into a homeostasis of self-renewal and differentiation. We utilize the developing *Drosophila* ovary as a versatile system to gain insights into the orchestration of cell type specific morphogenesis programs and the coordination of somatic gonad formation with primordial germ cell (PGC) development. A hormonal brain-to-gonad axis has been identified which coordinates gonadogenesis in *Drosophila*, similar to vertebrate systems (Dagklis et al., 2015; Gancz et al., 2011; Hodin and Riddiford, 1998; Sower et al., 2009). Here, distinct peaks of the steroid hormone ecdysone dictate the timing of somatic cell proliferation and maturation, and initiate PGC differentiation *via* a yet unknown mechanism. The exploration of this mechanism is the focus of this study.

The *Drosophila* gonads form at the end of embryogenesis when the progenitors of the germline, the PGCs, coalesce with the somatic gonadal precursor cells (Boyle and DiNardo, 1995; Gilboa and Lehmann, 2006; Li et al., 2003; Sano et al., 2012). During ovarian development, which starts in the first instar larva and continues throughout metamorphosis, PGCs divide and proliferate in coordination with the directly associated intermingled cells (ICs), while other somatic cell types diversify (Figure 1A) (Gilboa and Lehmann, 2006). An early ecdysone peak between early 3^rd^ larval instar (EL3) and mid 3^rd^ larval instar (ML3) (~90h ael) (Figure 1A) initiates the differentiation of the germline stem cell (GSC) niche, including the terminal filaments (TF) and cap cells (CC) (Gancz et al., 2011; Warren et al., 2006). The developing somatic niches associate with a fraction of PGCs and subsequently maintain them in an undifferentiated state through larval and pupal development and into adulthood (Gilboa and Lehmann, 2004; Song et al., 2004; Xie and Spradling, 1998, 2000). GSC maintenance relies on niche-secreted Bone Morphogenetic Protein (BMP) family ligands, such as Decapentaplegic (Dpp) and Glass bottom boat (Gbb) and the activation of Dpp/Gbb receptors in germ cells. Receptor-mediated phosphorylation of the Drosophila SMAD, Mothers against Dpp (Mad), blocks the germ cell differentiation pathway by binding to the promoter of the key differentiation factor *bag of marbles (bam)* (Chen and McKearin, 2003b; Song et al., 2004; Xie and Spradling, 1998, 2000). However, there are more PGCs than needed to fill all the niches. The remaining PGCs outside the niche experience only reduced levels of Dpp/Gbb signals, yet do not upregulate *bam* expression and do not initiate differentiation until the late 3^rd^ larval instar (LL3). Thus additional, still unknown regulators of *bam* transcription that act largely independent of Dpp/Gbb must mediate the temporal transition toward differentiation (Gancz et al., 2011). This switch towards PGC differentiation is temporally controlled by a late larval ecdysone pulse at ~100h ael preceding pupal development (Figure 1A) (Gancz et al., 2011; Warren et al., 2006).

**Figure 1.**
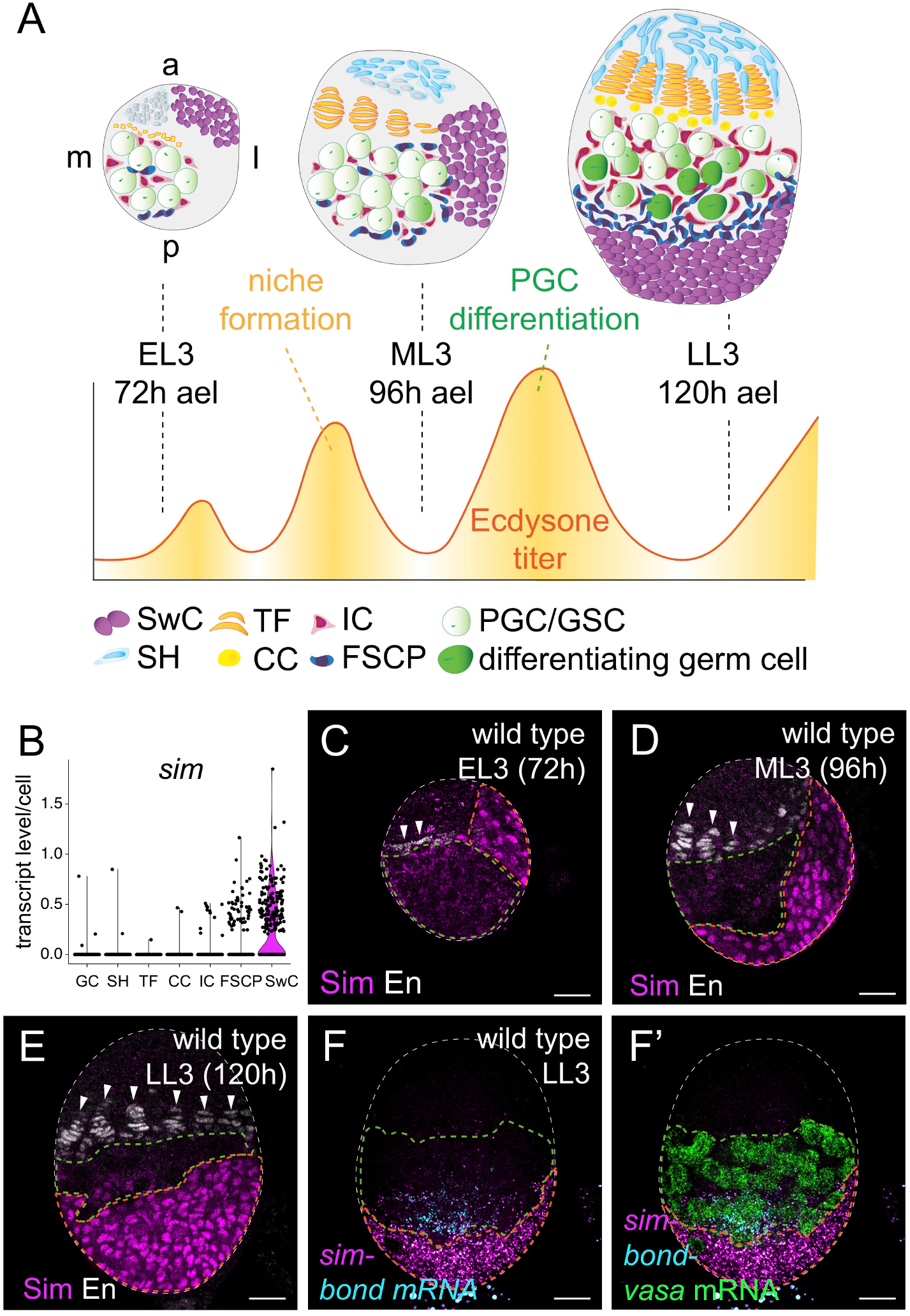
**Somatic swarm cell morphogenesis establishes a posterior gonad domain** (A) Schematic representation of larval ovary development. At EL3 (72h after egg lay (ael), primordial germ cells (PGCs in light green) are proliferating and in close contact with intermingled cells (ICs in red) and follicle stem cell precursors (FSCPs in dark blue). Terminal filament progenitors (TFs in orange) and sheath cells (SH in light blue) are specified at the anterior; swarm cells (SwCs in purple) are located antero-lateral. An early peak of the steroid hormone ecdysone between EL3 and ML3 (96h ael) induces niche formation as evidenced by TF cells exiting the cell cycle and organizing into stacks. By SH cells make up the anterior compartment, SwCs have moved to dorso-lateral positions. A late peak of ecdysone between ML3 and LL3 (120h ael) initiates PGC differentiation; germ cells close to the forming niches (TFs and cap cells (CC) in yellow) are maintained as germline stem cells (GSCs), while posterior PGCs (dark green) differentiate. SwCs form the posterior ovary compartment at LL3. (B) Violin plot from scRNA-seq. data for the SwC enriched gene *sim*; gene expression levels (y axis) for each ovarian cell cluster (x axis) are given, each dot represents a cell. (C-E) Anti-Sim antibody detects SwC compartment (orange outline) during development. Terminal filament morphogenesis (arrowheads) marked by anti-Engrailed (En) antibody (white) was used as independent, internal marker for developmental stages (EL3-LL3); germ cell compartment is outlined in green. (F) Spatial relationship of SwCs (*sim*) with respect to PGCs (*vasa*) and FSCPs (*bond*) detected by HCR *in-situ* hybridization at LL3. Scale bars indicate 10 μm.

It is unclear how PGC differentiation is controlled by ecdysone because PGCs do not express ecdysone receptors. It has therefore been proposed that PGCs interpret ecdysone pulses *via* a secondary cue relayed by somatic cells (Gancz et al., 2011). The identity of the somatic cell type that translates the ecdysone message into a trigger for PGC differentiation and the signaling pathway that transmits and integrates this message remain elusive.

We utilized a recently generated single cell RNA atlas of LL3 gonads (Slaidina et al., 2020) to search for a relay cell type and the elusive signal. We identified the somatic swarm cells (SwCs) as mediators of the ecdysone signal for posterior gonad formation and PGC differentiation. SwCs constitute a major cell type of the larval gonad that undergoes long distance morphogenetic movements from the anterior to the posterior of the gonad (Couderc et al., 2002). We show that ecdysone has a dual role in SwCs: the early ecdysone pulse initiates SwC morphogenesis and establishes a transitory signaling center. Once SwCs reach the posterior of the gonad, the late ecdysone peak induces expression of the perforin-like molecule Torso-like (Tsl) in SwCs. Tsl acts as a soma-to-germline signal that stimulates PGC differentiation *via* the Torso receptor tyrosine kinase (RTK). Activation of Torso RTK promotes PGC differentiation by releasing Krüppel-mediated transcriptional repression of *bam*. Intriguingly, SwC signaling function is limited to the larva to pupa transition, the only period during development when SwCs are in close proximity to the germline. Thus SwCs act as a transitory signaling center that aids in coupling general gonad morphogenesis with niche establishment and the initiation of PGC differentiation.

## RESULTS

### Swarm cell migration drives gonad morphogenesis and engages in transient interactions with PGCs

The *Drosophila* larval gonad at LL3 is comprised germ cells and of six somatic cell types, and the majority of somatic cells directly support germline development (Bolivar et al., 2006; Gilboa and Lehmann, 2004, 2006; Slaidina et al., 2020; Xie and Spradling, 2000). The somatic swarm cells (SwCs), also called basal cells, which generate the large posterior domain of the ovary, have not been functionally analyzed yet. Swarm cells originate from the anterior of the gonad and then move dorso-laterally, towards the posterior (Couderc et al., 2002). The timing of their migration and arrival at the posterior coincides with ecdysone peaks. As SwC migration proceeds in close proximity, initially past and then beyond, the location of the PGC population, we reasoned that the orchestrated migration of SwCs could act as a possible timer for germ cell differentiation (Figure 1A and 1C-E).

To analyze SwC behavior and function, we probed our recently generated single cell RNA sequencing (scRNA-seq.) data-set of LL3 ovaries for SwC gonadal cell type signature genes (Slaidina et al., 2020). We identified *single-minded* (*sim*) and *crossveinless-2* (*cv-2*) as highly and quite specifically expressed in SwCs (Figures 1B and S1A). Antibodies directed against Sim labeled SwC nuclei, and allowed us to observe them during gonad development (Figures 1C-E). To assess developmental stages accurately and independently of SwC morphogenesis, we used the progression of terminal filament formation labeled with anti-Engrailed (En) antibodies (Figure 1C-E) (Godt and Laski, 1995; Sahut-Barnola et al., 1995). SwCs can be readily detected anterolateral at EL3 and, at ML3, on the lateral and dorsal side of the gonad (Figure 1C-D). At LL3, most SwCs have completed their movements, forming a new posterior gonadal domain (Figure 1E) (Couderc et al., 2002). At this position they are in close proximity to the posteriorly located pool of PGCs and just below the follicle stem cell and follicle cell precursors (FSCPs) as shown by mRNA *in-situ* hybridizations for *sim* and the FSCP marker *bond* (Figure 1F-F’) (Slaidina et al., 2020).

To better characterize and genetically manipulate SwCs, we identified Gal4 lines for *sim* and *cv-2* that directed expression in SwCs. Both drivers express in SwCs as early as L2 (Figure 2A) and continued to be expressed during their migration (Figure 2B-D). *cv-2*-Gal4 had a somewhat broader expression domain than *sim*-Gal4 in SwCs, and also drove marker expression in some TFs, CCs and ICs (Figure S1B). Both these drivers largely recapitulate the expression pattern of the endogenous *sim* RNA, however, as noted before, their expression is sparse and not all SwCs are labelled (Slaidina et al., 2020). Overall, both drivers provide an excellent tool to analyze SwC behavior and function.

**Figure 2.**
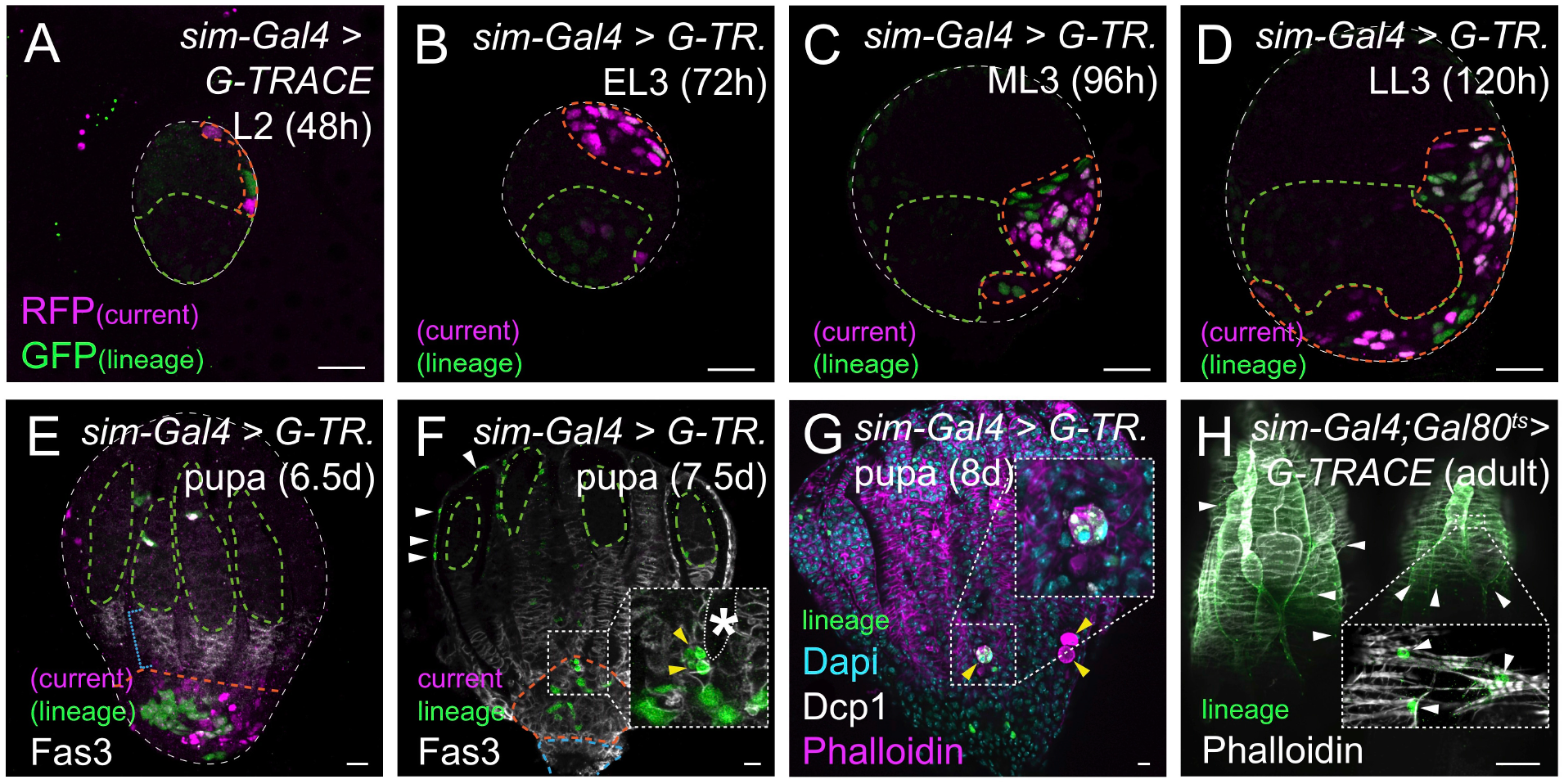
**Lineage tracing reveals transitory nature of swarm cells** (A-H) *sim*-Gal4 driven expression of a G-TRACE cassette follows SwC fate: current expression (RFP _current_) and lineage expression (GFP _lineage_); SwC locations and PGC positions are marked by orange and green outline, respectively. (A-G) G-TRACE labelling of current expression (RFP, magenta) reveals *sim* promoter activity in SwCs as early as L2 and throughout early pupa stages, recapitulating anterior to posterior SwC morphogenesis. Note that by 7.5d ael current expression can no longer be detected (A-H) G-TRACE labelling of lineage (GFP, green) reveals SwC fate. (E-F) In 6.5-7.5 d pupae SwC descendants are found at the very posterior end of gonad separated from germ cells (green outline) by follicle cells labeled with anti-Fasciclin3 (Fas3) antibodies (blue bracket). (F) By 7.5d the SwC cell lineage contributes to the calyx, which connects the ovary to the oviduct (blue outlined); few lineage-labeled cells can also be found in the developing peritoneal sheath (white arrowheads). (F-G) Rounded SwCs (yellow arrowheads in insets in F and G) at the base of developing ovarioles (extra-ovariolar cavity, white outline and asterisk in F) are undergoing apoptosis marked by antibodies against cleaved Dcp-1 (G). (H) G-TRACE expression was restricted to day 5 and 6 ael via Gal80^ts^ to specifically follow fate of late larval SwCs into the adult. Lineage expression in adults was detected prominently in the peritoneal sheath (Phalloidin, white arrowheads). Scale bar indicates 10 μm, except 100 μm in H.

To study the fate of SwCs from larval to pupal stages and into the adult, we made use of G-TRACE lineage tracing. Briefly, this method uses a temperature sensitive dual reporter system, where a tissue specific Gal4 drives (1) UAS-FLP recombinase to generate GFP marked clones for lineage labeling by FRT recombination at a defined developmental time and (2) a UAS- fluorescent reporter (RFP) to capture real-time expression patterns (Evans et al., 2009). During larval stages, constitutive expression of the G-TRACE cassette labeled the SwC lineage as early as L2 and throughout their migration (Figure 2A-D). During early pupal stages (6.5d AEL), SwCs were displaced from the vicinity of germ cells by the expanding layer of FSCPs, labeled by Fasciclin 3 (Fas3) staining (Figure 2E, bracket). At mid-pupal stages (day 7.5-8 ael), no current expression of G-TRACE could be detected, and SwC descendants were found to contribute to the calyx, a structure connecting the ovary to the oviduct (Figure 2F). Notably, SwC numbers seemed to diminish during pupal stages and cells undergoing programmed cell death were found at the base of developing ovarioles (Figures 2G and S1C). It had been suggested previously that the content of these dying cells contributes to the lumen separating individual ovarioles (King et al., 1968). In adults, few SwC descendants were detected in the peritoneal sheath (Figure 2H) (Slaidina et al., 2020), when G-TRACE expression was restricted to a brief developmental window from 120h to 144h ael. This suggests, that most of the SwC descendants are lost during pupal stages. In summary, direct observation and lineage tracing demonstrates that SwCs represent a transitory cell population that is in close vicinity to PGC. While, they constitute the most prominent cell population in the larval ovary (Slaidina et al., 2020), they largely disappear during pupal stages with some descendants contributing to the outer ovarian sheath in the adult.

### Ecdysone signaling promotes swarm cell migration

Our lineage tracing demonstrates that SwCs constitute a transitory cell type in the developing ovary that moves from the anterior tip of the ovary to the posterior during larval stages. The formation of the posterior ovary domain has been reported to depend on ecdysone receptor (EcR) signaling (Gancz et al., 2011). Here, a dominant negative form of EcR, EcRA^W650A^ (from here on EcR^DN^), which is insensitive to hormone but instead constitutively binds to EcR target genes and represses them, was expressed in all somatic cells. Expression of EcR^DN^, in all somatic gonadal cells via *tj-*Gal4 during the critical periods resulted in a block of PGC differentiation and impaired somatic gonad development and morphogenesis (Gancz et al., 2011): block of TF differentiation, failure in Cap cell formation, defects in IC intermixing with germ cells and a complete lack of the posterior compartment of the gonad (Figure S1D-E). We visualized SwCs under these conditions and found their migration to be dramatically affected; they reached the lateral side of the ovary, but failed to migrate towards the posterior. SwC numbers were also reduced under these conditions (Figure 3A-B). To investigate whether ecdysone directly regulated SwC morphogenesis, we expressed EcR^DN^ in SwCs specifically via *sim*-Gal4 and *cv-2*-Gal4. The resulting SwC morphogenesis defects were similar, albeit less pronounced (Figure 3C-D): the majority of SwCs was found at antero-lateral positions in the gonad and SwC numbers were reduced. The specificity of these drivers for SwC cells was further supported by the fact that other somatic tissues, which were affected after global expression of EcR^DN^, seemed unaffected as formation of TF stacks occurred normally and only few ovaries exhibited mild IC intermingling defects (3/20 gonads, compared to 0/16 gonads in control; Figure S1F-G).

**Figure 3.**
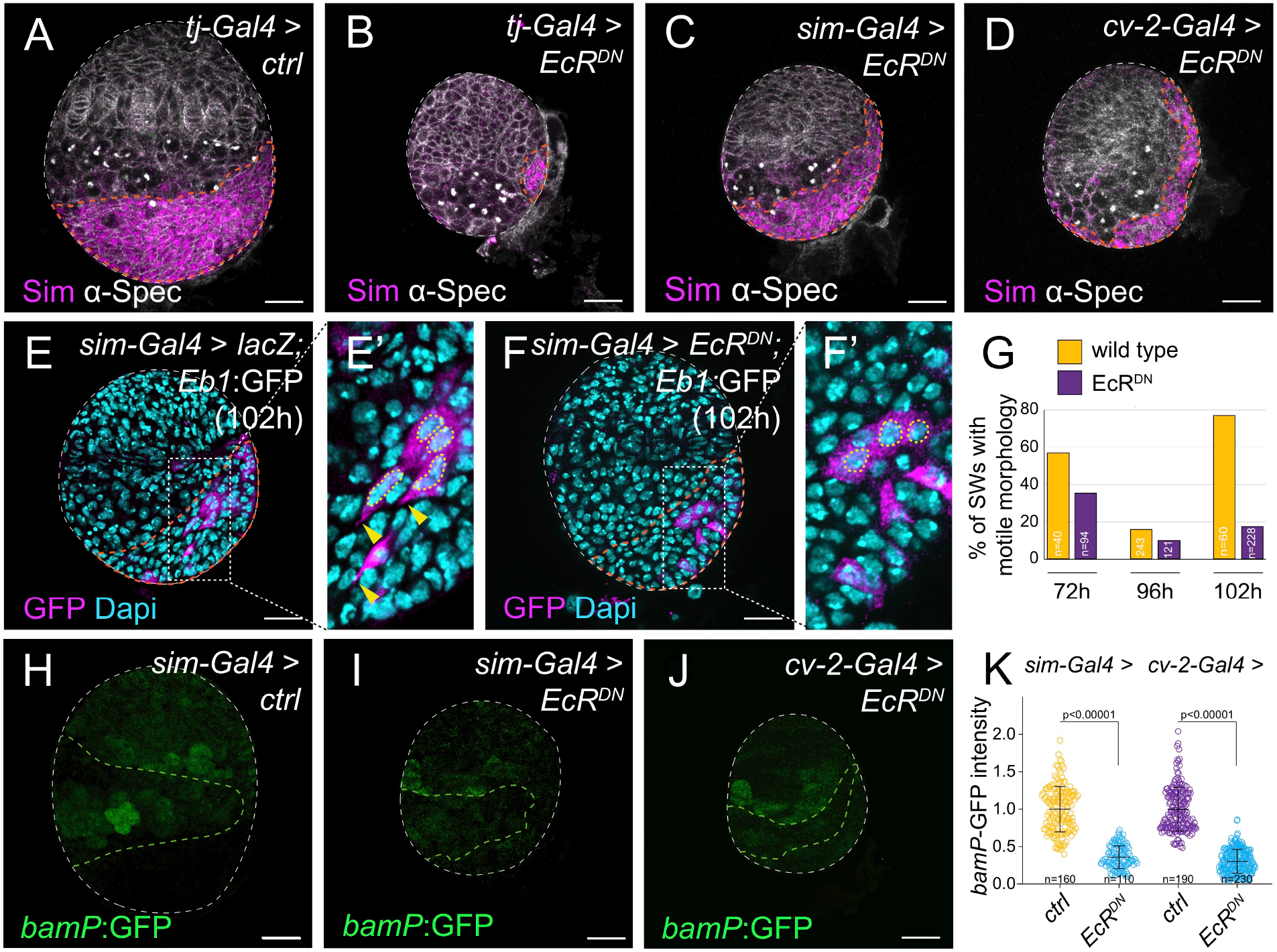
**Ecdysone pulses promote swarm cell morphogenesis and germ cell differentiation** (A-D) Effect of EcR^DN^ expression on SwC morphogenesis. SwCs are labeled with anti-Sim antibodies (orange outline); the outlines (plasma membranes) of all ovarian cells are marked by anti-α-Spectrin, which also recognizes the spectrosome, a round structure and hallmark of undifferentiated germ cells. (A) SwCs occupy the very posterior region in control ovaries at LL3 (120h ael); (B) pan-somatic (*tj-Gal4*)- or (C and D) SwCs specific (*sim-*Gal4, *cv-2*-Gal4) expression of EcR^DN^ impacts SwC number and morphogenesis. (E-F) Eb1:GFP and Dapi mark SwC cell protrusions and nuclear morphology, respectively. (E-E’) Wildtype SwCs initiate migration shortly after ML3, indicated by protrusions and elongated nuclear morphology in direction of movement (yellow arrowheads and outline) (Note that *sim-*Gal4 drives UAS expression in only a subset of SwCs). (F) SwCs expressing EcR^DN^ lack motile behavior. (G) Graphical representation of SwC motility index at EL3 (72h), ML3 (96h) and (>98h) in control and EcR^DN^ expressing SwCs. (H-J) PGC differentiation status is monitored by *bamP*:GFP, germ cell domain is outlined in green. (H) Wildtype PGCs initiate differentiation at LL3 stage; (I-J) *bamP*:GFP reporter expression is absent when EcR^DN^ is expressed in SwCs. (Note, the *bamP*:GFP reporter shows some ectopic expression in TFs, however, based on their position these cells are clearly distinct from the germ cell population.) (K) Graph depicting relative *bamP*:GFP expression levels with two independent SwC driver lines. Each data point represents a single PGC. Error bars represent std. Scale bars indicate 10 μm.

To investigate how ecdysone signaling impacts SwC motility we analyzed the morphology of wildtype and EcR^DN^ expressing SwCs at multiple developmental time-points by expression of a microtubule marker (Eb1:GFP). These studies defined three phases of SwC motility in the wild type: at EL3 (72h), wildtype SwCs exhibited clear signs of migratory behavior as judged by their elongated nuclear morphology and generation of microtubule-rich cellular protrusions towards the direction of movement (Figure 3E). These motility indicators were largely absent at ML3 (96 h), when SwCs reached a medio-lateral position, but resumed by 98h, and the vast majority of wildtype SwCs exhibited elongated nuclear morphologies and protrusions directed towards the posterior of the gonad at 102h (Figures 3E-G and S1H). EcR^DN^ expressing SwCs at EL3 displayed reduced motile characteristics compared to wild type (Figure 3G). Similar to wild type, EcR^DN^ expressing SwCs failed to display cell morphologies indicative of motile behavior at ML3 (96h), when SwCs apparently pause at the dorso-lateral side of the gonad (Figure 3G). However, subsequently, and in contrast to wild type, EcR^DN^ expressing SwCs did not show any migratory morphologies after ML3, (Figures 3E-G).

This suggest that discrete ecdysone pulses initiate SwC motility behavior. Considering the timeline of their migration, a pulse at L2 and possibly the early ecdysone cue at 90h stimulates SwC migration. A several hour delay between pulse and tissue response has been noted before (Ashburner, 1990; Regan et al., 2013; Stoiber et al., 2016), and results from the hierarchical expression cascade down-stream of ecdysone, where early response genes (e.g. transcription factors, chromatin remodelers) activate tissue specific transcriptional programs or expose additional binding sites for EcR (Uyehara and McKay, 2019; Yamanaka et al., 2013).

### Swarm cells relay Ecdysone signals to initiate timely PGC differentiation

Discrete pulses of ecdysone initiate the differentiation of the GSC niche (TFs and CCs) (early peak), including SwC migration, and the differentiation of PGCs (late peak) (Figure 1A and S1D-E) (Gancz et al., 2011). However, the somatic cell type that relays the later pulse to PGCs remains elusive. During their migration, SwCs are transiently in close proximity to PGCs and can extend protrusions towards them (Figure S1I). Further, the time of their arrival at the gonad posterior aligns well with the initiation of PGC differentiation. This suggests that SwCs may relay the late, prepupal peak of ecdysone that initiates PGC differentiation (Figure 1A) (Gancz et al., 2011). We therefore expressed EcR^DN^ specifically in SwCs and assessed PGC differentiation status with a transcriptional reporter for the key differentiation gene *bag of marbles* (*bamP*:GFP) (Chen and McKearin, 2003b). Under control conditions, *bamP*:GFP expression in PGCs was detected at LL3 (Figure 3H). Intriguingly, signal from the *bamP*:GFP reporter was barely detectable at LL3 when EcR^DN^ was expressed in SwCs *via sim*-Gal4 or *cv-2*-Gal4 (Figures 3I-K). These results suggest that EcR signaling in SwCs is required for proper timing of PGC differentiation, likely by inducing expression of a SwC to PGC signal.

To identify the secondary signaling molecules expressed by SwCs, we made use of the scRNA-seq. data set of LL3 larval gonads (Slaidina et al., 2020). Among the 5963 transcripts expressed in SwCs at LL3 (0.05 average expression cut off), only 130 were predicted to be present on cell surfaces or in the extracellular space by the PANTHER classification system (Mi et al., 2019). Of those, only one transcript was highly enriched in SwCs: the perforin-like molecule Torso-like (Tsl) (Figure 4A). RNAi mediated knockdown of *tsl* in SwCs resulted in diminished levels of *bamP*:GFP, whereas overexpression of *tsl* caused precocious PGC differentiation as evidenced by elevated *bamP*:GFP reporter levels and presence of branched fusomes, a hallmark of germ cell cysts normally present only at later stages (Figures 4B-E and S2A). These results suggest that Tsl acts as a novel soma to germline signal, regulating PGC differentiation status.

**Figure 4.**
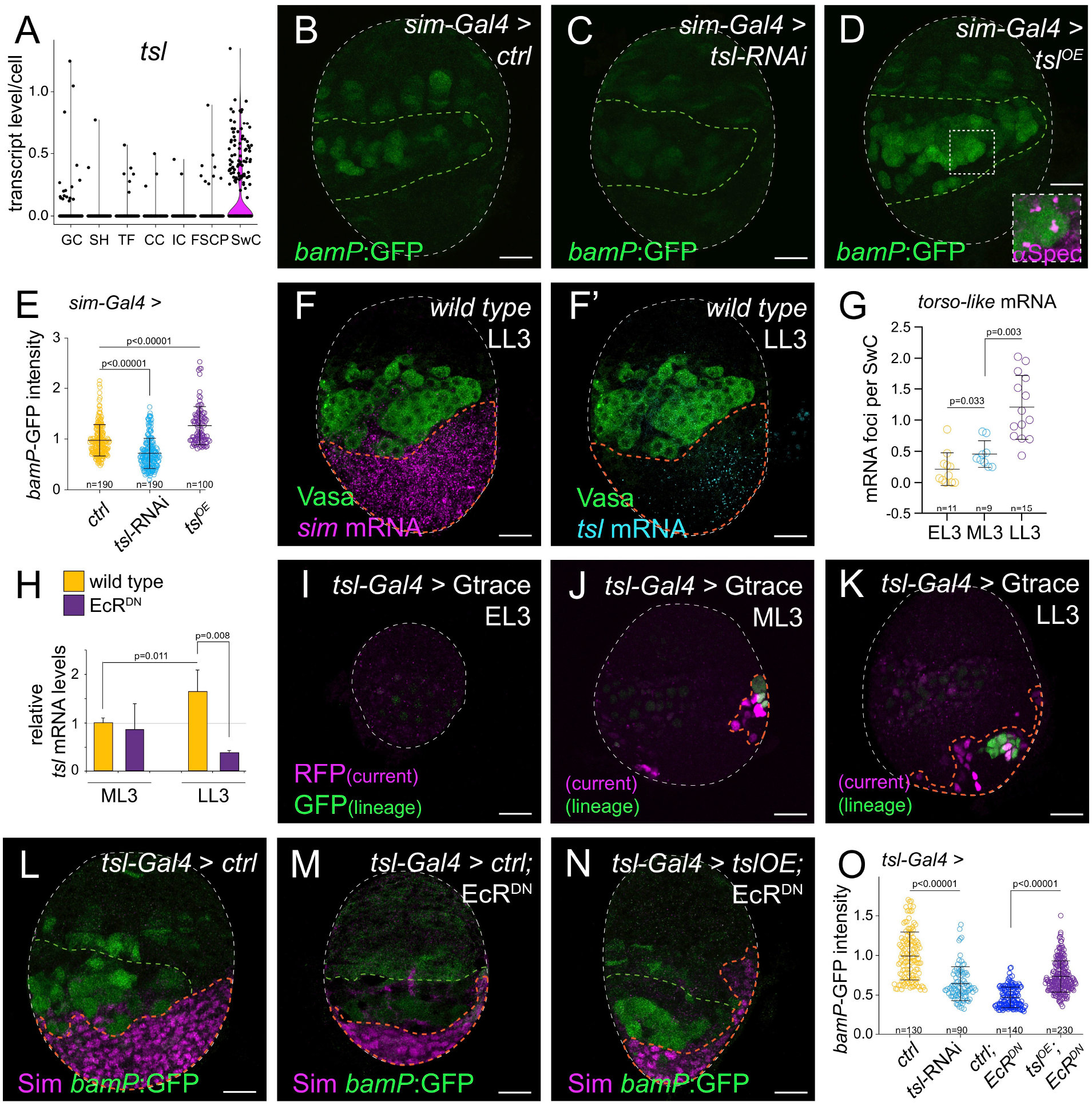
**Swarm cells express *tsl* to initiate PGC differentiation in response to a late larval ecdysone pulse** (A) Violin plot from scRNA-seq. data for the SwC enriched gene *torso-like (tsl)*; gene expression levels (y axis) for each ovarian cell cluster (x axis) are given, each dot represents a cell. (B-D) PGC differentiation status at LL3 is imaged by PGC differentiation marker *bamP*:GFP, germ cell domain is outlined in green. Compared to LL3 control (B), where some PGCs initiated differentiation as revealed by sparse *bamP*:GFP expression, in *tsl* RNAi knock-down (C) *bamP*:GFP expression was decreased, and in *tsl*^*OE*^ (D) *bamP*:GFP expression was increased. Precocious PGC differentiation is highlighted by presence of branched fusomes (anti-α-Spectrin, inset in D). (E) Graph quantifying *bamP*:GFP expression levels as exemplified in B-D. Each data point represents a single PGC. (F, F’) HCR *in*-*situ* hybridization for *tsl* and *sim* mRNAs at LL3, germ cells are labeled with anti-Vasa antibodies. (G) Graph showing number of *tsl* mRNA foci detected in SwCs at different developmental stages; each data-point represents a single ovary. (H) qPCR measurements showing an increase in *tsl* mRNA levels from ML3 to LL3 in wildtype, but not when EcR^DN^ was expressed in soma. (I-K) *tsl*-Gal4 driven expression of the G-TRACE cassette. No expression was detected at EL3 (I), few SwCs expressed G-TRACE at ML3 (J) and many were labeled at LL3 (K). (L-N) SwCs are labeled *via* anti-Sim antibodies (orange outline), PGC differentiation status is revealed by *bamP*:GFP (germ cell domain outlined in green). (L-M) *tsl*-Gal4 driven EcR^DN^: SwC migration was unaffected but PGC differentiation was delayed. (N) Reintroduction of *tsl* rescues the EcR^DN^ phenotype, placing Tsl function downstream of EcR^DN^. Note the effect of EcR^DN^ on SwC number is independent of *tsl* expression. (O) Graph depicting relative *bamP*:GFP expression levels as exemplified in L-M. Each data point represents a single PGC. Error bars represent std. Scale bar indicates 10 μm.

To determine whether Tsl functions as a temporal switch for PGC differentiation, we analyzed *tsl* mRNA levels at EL3, ML3 and LL3 using a highly sensitive RNA *in-situ* hybridization chain reaction (HCR) (Choi et al., 2018). While, *tsl* mRNA was barely detectable in SwCs at EL3 and ML3, significantly more *tsl* mRNA foci could be detected per SwC at LL3 (Figures 4F-G, S2B-D), suggesting developmental upregulation of *tsl* expression at LL3. *tsl* mRNA was also detected in few FSCPs, ICs and germ cells at LL3, consistent with the scRNA seq. data. The temporal regulation of *tsl* expression was recapitulated by a previously generated *tsl* promoter fusion construct (*tsl-*Gal4) (Grillo et al., 2012). Using the lineage tracing G-TRACE method, we found the UAS-RFP reporter expressed sparsely in SwCs at ML3, while a much larger fraction of SwCs was labeled at LL3 (Figures 4I-K). Finally, we directly compared *tsl* mRNA levels by qPCR from isolated ovaries and found that *tsl* expression was increased two-fold between ML3 to LL3. Importantly, this up-regulation was not detected in EcR^DN^ ovaries, suggesting that initiation of *tsl* expression occurs downstream of ecdysone (Figure 4H). Together, these results suggest that the late ecdysone peak initiates *tsl* expression in SwCs, which promotes timely PGC differentiation.

To separate ecdysone’s early role in SwC migration from a possible later role in relaying a differentiation signal to PGCs, we made use of the expression pattern of the *tsl*-Gal4 driver to block EcR signaling in SwCs. Indeed, late expression of EcR^DN^ via *tsl*-Gal4 did not affect SwC motility and SwCs did form a clear posterior domain by LL3, confirming that SwC migration is initiated by the early cue (Figure 4L-M). SwC numbers were reduced compared to wild type suggesting a continuous requirement of ecdysone for SwC proliferation. Importantly, PGCs did not initiate differentiation at LL3 under these conditions, indicating that the late ecdysone peak promotes PGC differentiation via Tsl (Figures 4L-O). In support, *tsl*-Gal4 driven expression of *tsl*-RNAi in SwCs blocked PGC differentiation at LL3. *tsl* overexpression in EcR^DN^ SwCs rescued PGC differentiation defects supporting a model whereby Tsl functions downstream of EcR in PGC differentiation (Figures 4M-O). Thus, ecdysone-dependent expression of Tsl in SwCs acts as a soma-to-germline communicator that is both necessary and sufficient to promote timely PGC differentiation.

### Somatic Tsl promotes Torso signaling in PGCs to initiate their timely differentiation

To determine whether Tsl may be directly acting on PGCs and how it is integrated into the PGC differentiation pathway, we explored the signaling cascade downstream of Tsl (Casanova et al., 1995; Johnson et al., 2015). In the fly embryo, Tsl facilitates the activation of Torso by its ligand Trunk (Trk). While the exact mechanisms by which Tsl acts are still uncertain, the pathway downstream of Torso RTK is well defined and leads via the MAP kinase signaling cascade to the activation of the transcription factors tailless (*tll*) and *huckebein* (*hkb*); these, in turn, instruct the development of terminal structures at the very anterior and posterior ends of the embryo (reviewed in (Goyal et al., 2018; Mineo et al., 2018). Consistent with the hypothesis that the complete Torso RTK signaling cascade functions in PGCs (Figure 5A), we detected *torso* and *trk* transcripts specifically in PGCs at LL3 in our scRNA-seq. dataset (Figure S2E-F) as well as by HCR *in-situ* hybridization (Figure 5B) and qPCR of LL3 and ML3 gonads (Figure S2G). We could also detected the transcript of the Torso pathway effector *tailless* (*tll*) in PGCs by HCR (Figure 5B).

**Figure 5.**
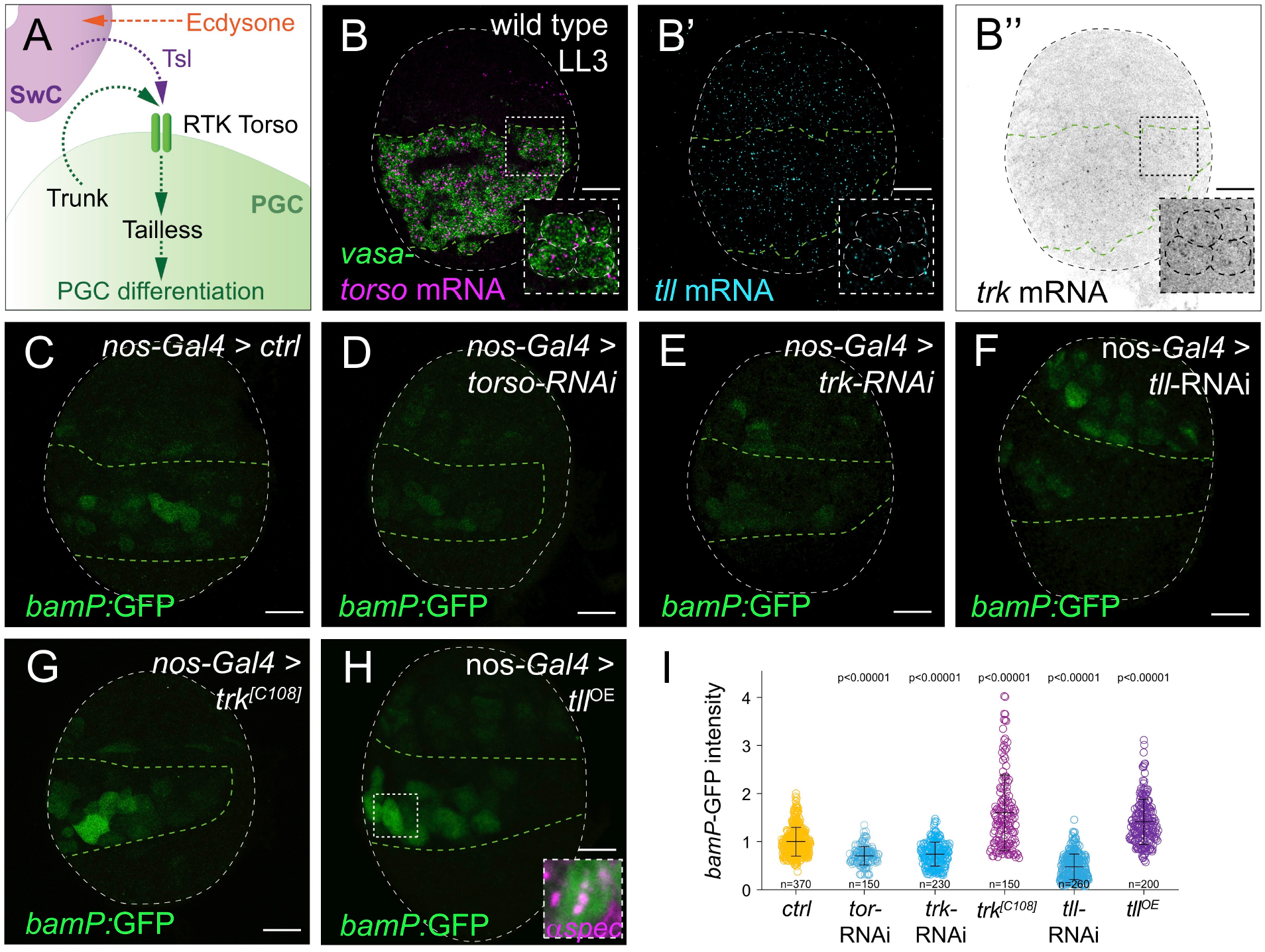
**Temporal control of Tsl expression in Swarm cells promotes PGCs differentiation via RTK Torso signaling** (A) Schematic summary of SwC-to-germline signaling via Tsl-Torso pathway. (B) HCR *in-situ* hybridizations for *torso*, *trunk* (*trk)*, *tailless* (*tll*) and *vasa* mRNAs. (C-H) PGC differentiation status is indicated by *bamP*:GFP, germ cell domain is outlined in green. RNAi-mediated knock-down of *torso* (D), *trk* (E), or *tll* (F), resulted in decreased *bamP*:GFP expression when compared to control. Expression of an activated version of Trk (*trk^[C108]^*) (G), and overexpression of *tll* (H), resulted in precocious PGC differentiation. (I) Graph depicting relative *bamP*:GFP expression levels as exemplified in C-H. Each data point represents a single PGC; P values in relation to control. Error bars represent std. Scale bar indicates 10 μm.

To test the involvement of this signaling cascade in this new context, we performed RNAi knockdown experiments using the germline specific driver *nanos*-Gal4. Knockdown of either *torso*, *trk* or *tll* resulted in decreased *bamP*:GFP levels compared to wild type consistent with a role in PGC differentiation (Figures 5C-F). Furthermore, expression of an activated version of the ligand Trk (*trk^C108^*) as well as overexpression of the downstream target *tll* resulted in precocious PGC differentiation, as early as ML3 (Figures 5G-I and S2H-I). These results demonstrate that activation of Torso RTK signaling cascade temporally controls PGC differentiation.

### Torso signaling activates key PGC differentiation factor *bam* by relieving Krüppel mediated repression

Our results show that PGC differentiation is temporally regulated by ecdysone induced expression of Tsl in SwCs, which triggers activation of the RTK Torso signaling cascade in PGCs. However, it remained unclear how Torso pathway activation is integrated into the PGC differentiation pathway ultimately leading to the expression of *bam*. During the ML3 to LL3 transition only a fraction of PGCs are incorporated into newly formed niches and are subsequently maintained as GSCs by niche secreted Dpp. In contrast to germ cells close to the niche, PGCs outside the niche experience only reduced levels of Dpp signaling, as indicated by absence of pMad in PGCs (Gancz et al., 2011; Zhu and Xie, 2003). Thus *bam* is not transcriptionally repressed by niche signals, yet, PGCs require an ecdysone input to express *bam* and thus initiate differentiation only at a later time point. This observation implicates the existence of additional, temporally controlled repressors of *bam* transcription that acted independently of Dpp. We reasoned that activation of the Torso-Tll cascade at LL3 might relieve this repression. To identify potential repressors, we asked whether targets of the Torso pathway known for their role as repressor in during embryogenesis are expressed in larval PGCs. We found that the repressor gene Krüppel (Kr) is expressed in PGCs at LL3 albeit at very low levels (Gaul and Jackle, 1987; Slaidina et al., 2020; Steingrimsson et al., 1991). PGC specific *kr* knockdown by RNAi transgenes caused precocious PGC differentiation (Figures 6A-D and S3A-B). Consistent with Kr acting as a repressor of PGC differentiation outside of the niche domain, PGCs in close proximity to the somatic niche were unaffected by expression of *kr*-RNAi constructs as judged by lack of *bam*:GFP expression and normal levels of pMad in these cells (Figures 6B and S3C-D). Conversely, overexpression of Kr in PGCs effectively blocked *bamP*:GFP reporter expression (Figure 6C-D), suggesting Kr function represses the differentiation program in PGCs from early larval stages on.

**Figure 6.**
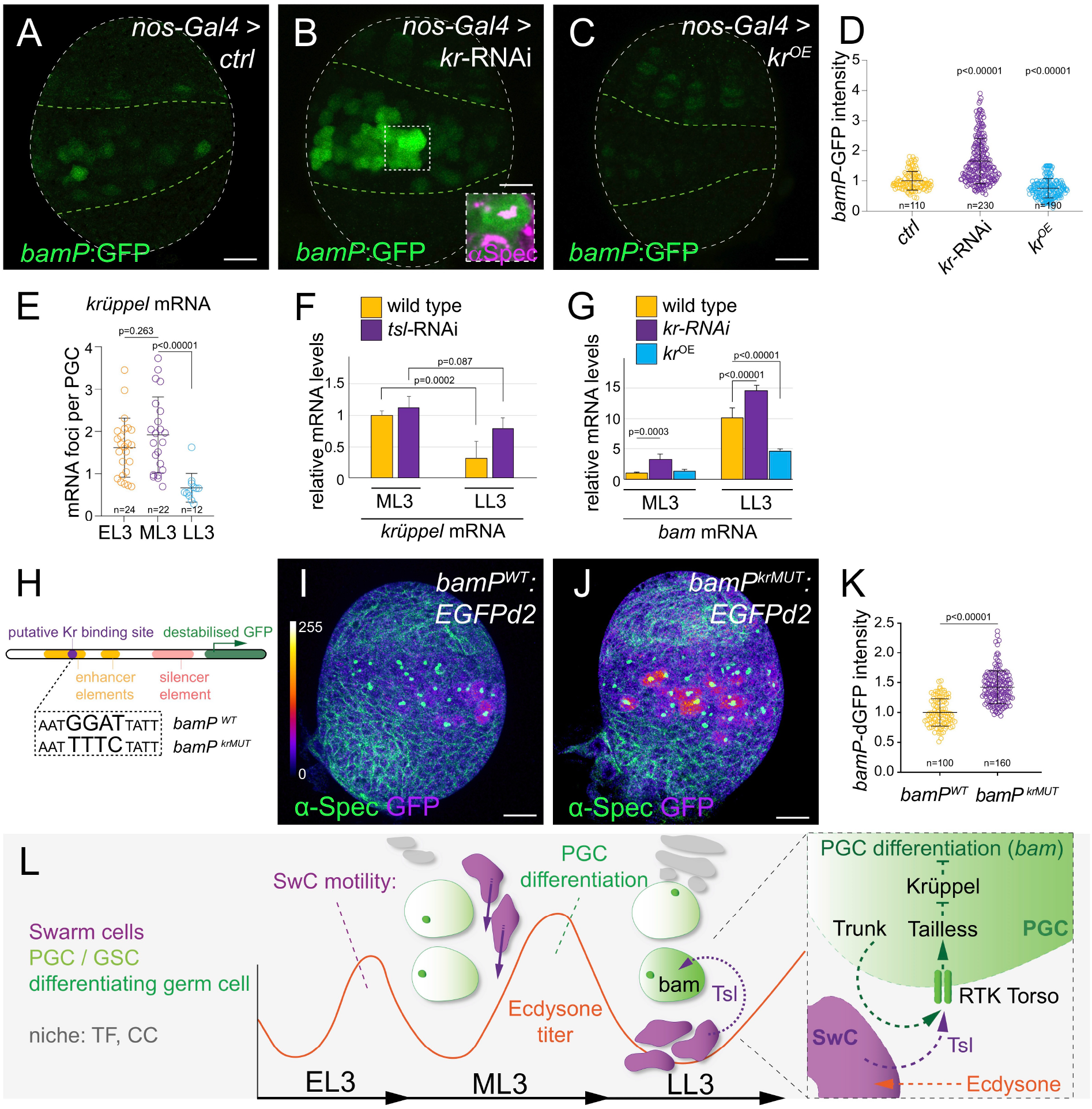
**Torso pathway promotes PGCs differentiation by alleviating Krüppel-mediated repression of *bam* promoter activity.** (A-C) *bamP*:GFP expression indicates differentiation status in LL3 gonads, germ cell domain is outlined in green. (B) PGC specific *kr*-RNAi knockdown results in precocious differentiation, (C) *kr*^*OE*^ blocks differentiation. (D) Graph depicting relative *bamP*:GFP expression levels as shown in A-C. Each data point represents a single PGC; P values in relation to control. (E) Measurements of *kr* mRNA foci detected by HCR in PGCs at different developmental stages; each data-point represents a single ovary. (F) qPCR measurements show decrease of *kr* mRNA levels from ML3 to LL3 in wild type gonads but not when *tsl* RNAi was expressed in soma. (G) qPCR measurement of *bam* mRNA levels, showing significant increase from ML3 to LL3 in wildtype. *kr*-RNAi in germ cells resulted in elevated *bam* levels at ML3 while *kr* overexpression dampened *bam* transcription at LL3. (H) Schematic representation of reporter constructs for *bam* promoter activity. In *bamP^wt^*:EGFPd2, the *bam* core promoter sequence was fused to a destabilized EGFP to report changes in promoter activity; known enhancer elements (orange) and the Mad/Med targeted silencer element (red) are indicated (Chen and McKearin, 2003a; Chen and McKearin, 2003b). In *bamP^Krmut^*:EGFPd2A a putative Kr binding site (purple) was mutated. (I-J) Reporter expression is increased when the Kr binding site is mutated. Note that germ cells in contact to niche do not express *bamP^Krmut^*:EGFPd2A. (K) Graph depicting relative expression levels of the *bam* transcriptional reporters as exemplified in I-J. Each data point represents a single PGC; Error bars represent std. Scale bar indicates 10 μm. (L) Model for Ecdysone control on SwC morphogenesis and PGC signaling: an early ecdysone peak initiates differentiation of somatic tissues and promotes posterior movement of SwCs. A second ecdysone peak, directly perceived by SwCs, evokes soma-to-germline signaling: Timed Tsl expression in SwCs activates RTK Torso in PGCs and initiates a downstream signaling cascade that relieves Kr-mediated transcriptional repression of *bag of marbles*, allowing for PGC differentiation.

We next investigated whether and how Kr levels are developmentally regulated to allow initiation of PGC differentiation at LL3. HCR *in-situ* hybridizations at EL3, ML3 and LL3 showed that *kr* mRNA is expressed in PGCs at early stages and is down-regulated by LL3 (Figures 6E and S3E-G), which coincides with the upregulation of *tsl* expression in SwCs. To test whether initiation of Torso signaling cascade represses *kr* transcription, we compared *kr* mRNA levels by qPCR in the presence and absence of Tsl. We found that *kr* mRNA levels remained high at LL3 in *tsl* knock-down condition, suggesting that *kr* is indeed regulated by the Tsl-Torso-Tll axis (Figure 6F).

Next, we asked whether the effects of Kr on PGC differentiation are mediated by changes in the transcriptional activity of the main differentiation gene *bam*. We performed qPCR experiments on whole ovaries at ML3 and LL3. In wild type, *bam* mRNA levels were upregulated 10 fold from ML3 to LL3, consistent with the initiation of PGC differentiation (Figure 6G). Conversely, upregulation of *bam* transcription was significantly blocked in PGCs overexpressing Kr, while PGCs expressing RNAi constructs against *kr* showed an increase in *bam*-mRNA levels already at ML3 compared to wild type (Figure 6G). These results indicated that Kr blocks early PGC differentiation by repressing *bam* transcription, possibly by directly controlling the *bam* promoter as has been shown for the Dpp downstream effector Smad (Chen and McKearin, 2003a; Chen and McKearin, 2003b). *In-silico* predictions of transcription factor binding sites in the *bam* promoter revealed a prominent near consensus-binding site for Kr that lies within the promoter region shown to be a critical enhancer element for *bam* transcription (Figure 6H) (Chen and McKearin, 2003b; Ho et al., 2009; Li et al., 2008). To test whether this putative Kr binding site functions to regulate *bam* expression we generated transcriptional reporters carrying either the wildtype *bam* core promoter sequences (*bamP*^wt^) or a promoter version with mutations in the predicted Kr binding site (*bamP*^krMUT^) (Figure 6H). These reporters were fused to a destabilized version of EGFP, which has a predicted half-life of two hours and thus provides a good estimate of *bam* transcriptional activity. We detected a significant increase in reporter expression when Kr binding sites are mutated (*bamP*^krMUT^) compared to the wildtype construct at LL3, suggesting precocious activation of *bamP* driven transcription (Figures 6I-K). Together our results suggest that Kr is both necessary and sufficient to block PGC differentiation via its role as a repressor of *bam* expression. As for *kr* RNAi knockdown, *bam* promoter de-repression was only observed in germ cells not associated with the niche, suggesting that the Dpp-Smad pathway can still exert its effect on this promoter fragment and acts independently of the Torso-Kr axis.

## DISCUSSION

Temporal and spatial coordination of cell proliferation, fate specification and tissue morphogenesis are hallmarks of organogenesis. This interplay between tissues is particularly important during development when stem cells are launched within their niches. During this critical period, developing niches may not be fully functional, yet stem cell progenitors have to remain in an undifferentiated state. Our results reveal another aspect of stem cell progenitor control, whereby the differentiation status of cells outside the niche has to be temporally aligned with organogenesis. Our studies identify the somatic swarm cells (SwCs) as a critical ecdysone target that transmits a soma to germline differentiation signal. An early ecdysone pulse initiates morphogenetic movements of SwCs from the anterior towards the posterior of the larval gonad. This new position brings SwCs into close contact with PGCs and, in response to a second ecdysone pulse, SwCs then transmit a differentiation signal to PGCs. This signal activates Torso receptor tyrosine kinase signaling in PGCs, which relieves repression by the transcriptional repressor Krüppel in PGCs, allowing for expression of the key germ cell differentiation gene *bam* (Figure 6L).

### Hormonal control of gonadogenesis

Steroid-mediated coordination of developmental processes is widely conserved. Pulses of steroid hormones evoke cell type and stage specific outcomes depending on their spatio-temporal regulation, titer and co-factors. In *Drosophila* gonadogenesis, the steroid hormone ecdysone serves a dual role: initiation of important early aspects of gonad morphogenesis, such as formation of the somatic niches to incorporate germline stem cells (GSC), SwC morphogenesis to establish a posterior gonadal compartment, and initiation of PGC differentiation indirectly via SwCs ((Gancz et al., 2011); this study).

Akin to ecdysone, in *C. elegans*, the steroid hormone dafrachronic acid (DA) regulates proper gonad morphogenesis, by ensuring correct migration of gonadal leader cells (DTC and LC), which elongate the gonadal arms (Antebi et al., 1998; Motola et al., 2006). DA further promotes germ cell proliferation indirectly at larval stages and has been shown to directly block proliferation of adult germ cells, suggesting a stage specific role of DA in soma and germline development (Mukherjee et al., 2017; Narbonne and Roy, 2006). In mammals, steroidogenic lineages develop only later in development (Herbison, 2016), and the contribution of exogenous signals to early gonadogenesis is less well understood. Analogous to ecdysone, Retinoid acid (RA) acts in a concentration dependent manner and binds nuclear receptors that regulate transcriptional programs in germ cells and somatic gonadal cells. In mice, RA is produced in the mesonephros and promotes timely meiotic entry of ovarian germ cells early in development (Bowles et al., 2016; Bowles et al., 2006; Chassot et al., 2020; Koubova et al., 2006; Vernet et al., 2020). Similar to the dual role of ecdysone, RA also regulates somatic gonadal support cell development, the extend of which is under investigation (Bowles et al., 2018; Minkina et al., 2017). In Humans, RA directly stimulates germ cell proliferation, differentiation and initiates meiosis of ovarian germ cells (Childs et al., 2011; Le Bouffant et al., 2010). These examples highlight a conserved function of systemic pulsatile cues in gonadogenesis. They act as temporal coordinates and orchestrate somatic gonad formation and germline development. This is in part achieved by promoting expression of secondary cell-cell communicators, which instruct surrounding cells as shown in this study. It remains to be determined how widespread cell-cell signaling in response to steroids/RA is employed in coordinating tissue development, and if similar pathways are involved.

While our data suggests that ecdysone signaling in SwCs is both necessary and sufficient to promote PGC differentiation, ICs and FSCPs may also contribute to this process. ICs are in close contact with PGCs, regulate PGC proliferation rates and adult IC descendants, the Escort cells support germ cell differentiation (Banisch et al., 2017; Gilboa and Lehmann, 2006; Kirilly et al., 2011; Schulz et al., 2002); and the FSCP lineage promotes proper oogenesis later in development (Slaidina et al., 2020). Both ICs and FSCPs express ecdysone receptors (Gancz et al., 2011; Slaidina et al., 2020), and *tsl* expression can be detected in few ICs and FSCPs at LL3 (Figures 4A and S2D).

### A transient signaling center

Transient signaling centers have been well described in many contexts. Known as signaling or organizing centers, discrete, specialized groups of cells serve as spatio-temporal point sources for signals that coordinate or evoke developmental processes. These cells, mostly transient in nature, undergo programmed cell death or incorporate into other tissues following their signaling function (Anderson and Stern, 2016; Basson, 2012). We have identified the somatic SwCs as a transitory cell type with signaling capacity in the developing ovary. SwCs play a dual role in ovary development: 1) a structural role, where SwC morphogenesis movements establish the posterior domain of the gonad (Couderc et al., 2002) and likely aid in connecting the ovaries to the oviduct; and 2) a signaling role to promote timely PGC differentiation.

The existence of transitory cell types during gonadogenesis has also been reported in *C. elegans*. The linker cell in *C. elegans* males migrates to elongate the gonadal arm, connects the reproductive tract to the exterior, and undergoes programmed cell death. However, no signaling function has been attributed to the linker cell (Antebi et al., 1997; Hedgecock et al., 1987; Kimble and Hirsh, 1979). In *C. elegans* hermaphrodites, the transitory anchor cell has pronounced patterning functions. It induces nearby epidermal precursor cells to generate vulval cells *via* an epidermal growth factor signaling pathway, patterns the uterus, and helps establish the physical connection of the epidermis with the uterus (Hill and Sternberg, 1992) (Newman et al., 1995, 1996).

In the case of SwCs, it remains to be determined whether they serve as a major signaling hub by relaying signals to other cell types in addition to PGCs. Observations of SwC migration and terminal filament maturation indicate that these movements are coordinated (Figure 1C-E): TF stack formation occurs in a medial to lateral fashion with the least mature TF stack in the vicinity of SwCs (Godt and Laski, 1995; Sahut-Barnola et al., 1996), which provides a spatial segregation of both processes. This coordination may involve SwC signals.

### Regulatory hierarchies for stem cell maintenance and differentiation

We identified a new signaling module in PGCs that regulates the timing of PGC differentiation. Our findings suggest that Kr functions in parallel to the well-established Dpp/pMad mediated signaling pathway throughout larval development (Gilboa and Lehmann, 2004; Song et al., 2004; Xie and Spradling, 1998, 2000). Both pathways block precocious PGC differentiation, however their actions are regulated in time and space. Before ML3, Dpp-mediated signaling and Kr repress PGC differentiation throughout the PGC compartment; from ML3 (96h) the reach of DPP becomes progressively restricted to the proximity of the newly formed niches to maintain a fraction of PGCs as GSCs for adulthood (Gancz and Gilboa, 2013; Gilboa and Lehmann, 2004; Song et al., 2002; Zhu and Xie, 2003). Precocious differentiation of PGCs outside the niches, which are not exposed to Dpp, is continuously repressed by Kr until mid-LL3 (~108h) ael when the Torso RTK pathway is activated and *bam* expression is first detected in PGCs (Gancz et al., 2011).

This dual repression system keeps PGCs in an uncommitted state to accommodate the complexity of somatic gonad morphogenesis. This allows terminal filament and cap cells to differentiate and form a functional niche, and allows FSCP positioning and specification before initiating the first round of germline cyst formation at LL3 (Gilboa, 2015; Slaidina et al., 2020). Any failures in this temporal and physical coordination is detrimental to fecundity as precocious PGC differentiation decreases the pool of future GSCs and precociously formed cysts likely undergo apoptosis (Gancz et al., 2011; Wang and Lin, 2004).

This dual repression system is integrated at the level of the *bam* promoter. A Kr binding site, responsive to Torso signaling, is located within a critical enhancer element of the *bam* promoter (Chen and McKearin, 2003a; Chen and McKearin, 2003b). Removal of this element results in low *bam* expression, suggesting Kr gates *bam* expression and does not fully abrogate expression as exerted by pMad/Medea (Chen and McKearin, 2003a; Chen and McKearin, 2003b). In line with the suggested mode of Kr function as a local quenching factor for nearby activators (Gray and Levine, 1996), we hypothesize that Kr interaction with this enhancer element blocks access to the *bam* promoter for still elusive transcriptional activators.

## Supplemental Figure legends

**Figure S1.**
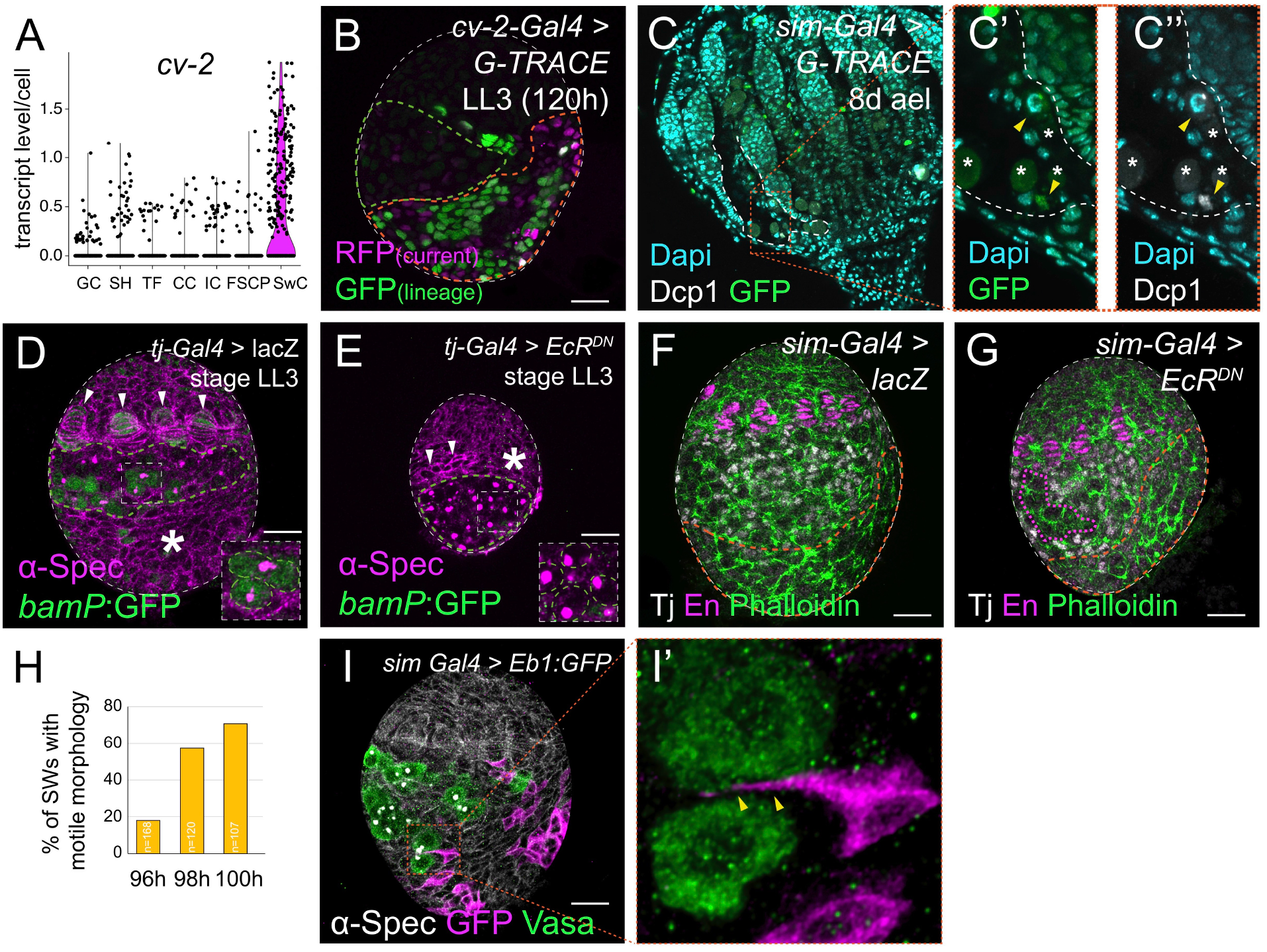
**(complementary to Figures 1, 2 and 3) Characterization of SwC driver lines and SwC fate** (A) Related to Figure 1B. Violin plot from scRNA-seq. data highlighting expression of the SwC enriched gene *cv-2*; gene expression levels (y axis) in each ovarian cell cluster (x axis) are given, each dot represents a cell. (B) Related to Figure 2D. *cv-2*:Gal4 driven expression of the G-TRACE cassette. Current expression (RFP_current_) and lineage expression (GFP_lineage_) at LL3; SwC cell location (orange outlined) and PGC position (green outline) are given. Note the pronounced reduction in current expression levels compared to lineage. (C) Related to Figure 2F-G. *sim*-Gal4 driven expression of G-TRACE at pupal stages. SwCs undergo apoptosis as indicated by anti-cleaved Dcp-1 antibody staining (arrowheads) and by advanced stage cell death morphology (asterisks) at the base of developing ovarioles in the extra-ovariolar cavity (white outline). (F-G) Related to Figure 3C-D. Besides SwC specific defects, somatic gonadal development is largely unaffected when EcR^DN^ is expressed specifically in SwCs with *sim*-Gal4. Cell outlines are labeled with Phalloidin, TFs with anti-Engrailed antibodies (En) and ICs with anti-Tj antibodies. SwC region is orange outlined. (F) Wild type ovary; (G) shows a mild IC intermingling defect in EcR^DN^ (magenta outline), together with SwC migration defects and slightly reduced gonad size. (H) Related to Figure 3E-G. Graphical representation of wild type SwC motility index at 96h, 98h and 100h, showing SwCs initiate migration shortly after ML3 (96h). (I-I’) Eb1:GFP marked cellular protrusions of SwCs (arrowheads) can probe in between germ cells (anti-Vasa antibodies).

**Figure S2.**
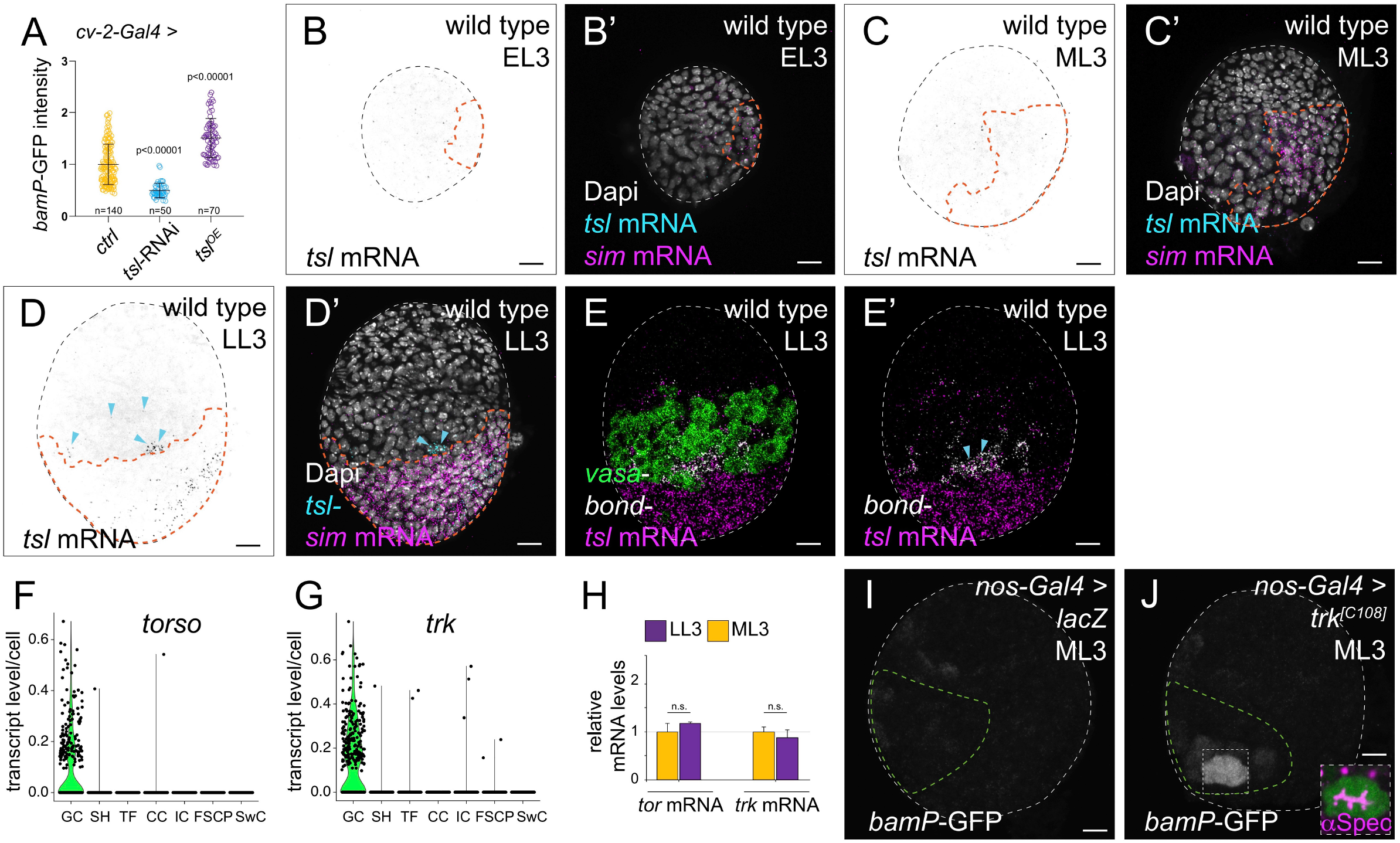
**(complementary to Figures 4 and 5) Torso signaling pathway functions to initiate PGC differentiation** (A) Related to Figure 4E. Graph depicting relative *bamP*:GFP expression levels comparing control to SwC specific (*cv-2*-Gal4) expression of *tsl*-RNAi- or *tsl*^*OE*^ constructs. Each data point represents a single PGC. P values in relation to control. (B-D; B’-D’) Related to Figure 4G. HCR *in situ* hybridization for *tsl* mRNA at different developmental stages. *sim* RNA expression pattern used as SwC cell marker (orange outline), Dapi labels all nuclei. Note *tsl* expression in few cells at FSCP position and in ICs (cyan arrowheads). (E-F) Related to Figure 5A-B Violin plots from scRNA-seq. data for the germ cells enriched genes *torso* and *trunk (trk); gene* expression levels (y axis) for each ovarian cell cluster (x axis) are given, each dot represents a cell. (G) Related to Figure 5A-B. qPCR measurements of *tor* and *trk* mRNA levels reveal no significant change in mRNA levels between ML3 and LL3. (H-I) Related to Figure 5G. Expression of *trk*^*[C108]*^ in PGCs results in precocious differentiation at ML3. PGC differentiation is marked by *bamP*:GFP expression and appearance of branched fusome, a morphological indicator of germ cell differentiation, visualized by anti α-Spectrin antibodies. Scale bars indicate 10 μm.

**Figure S3.**
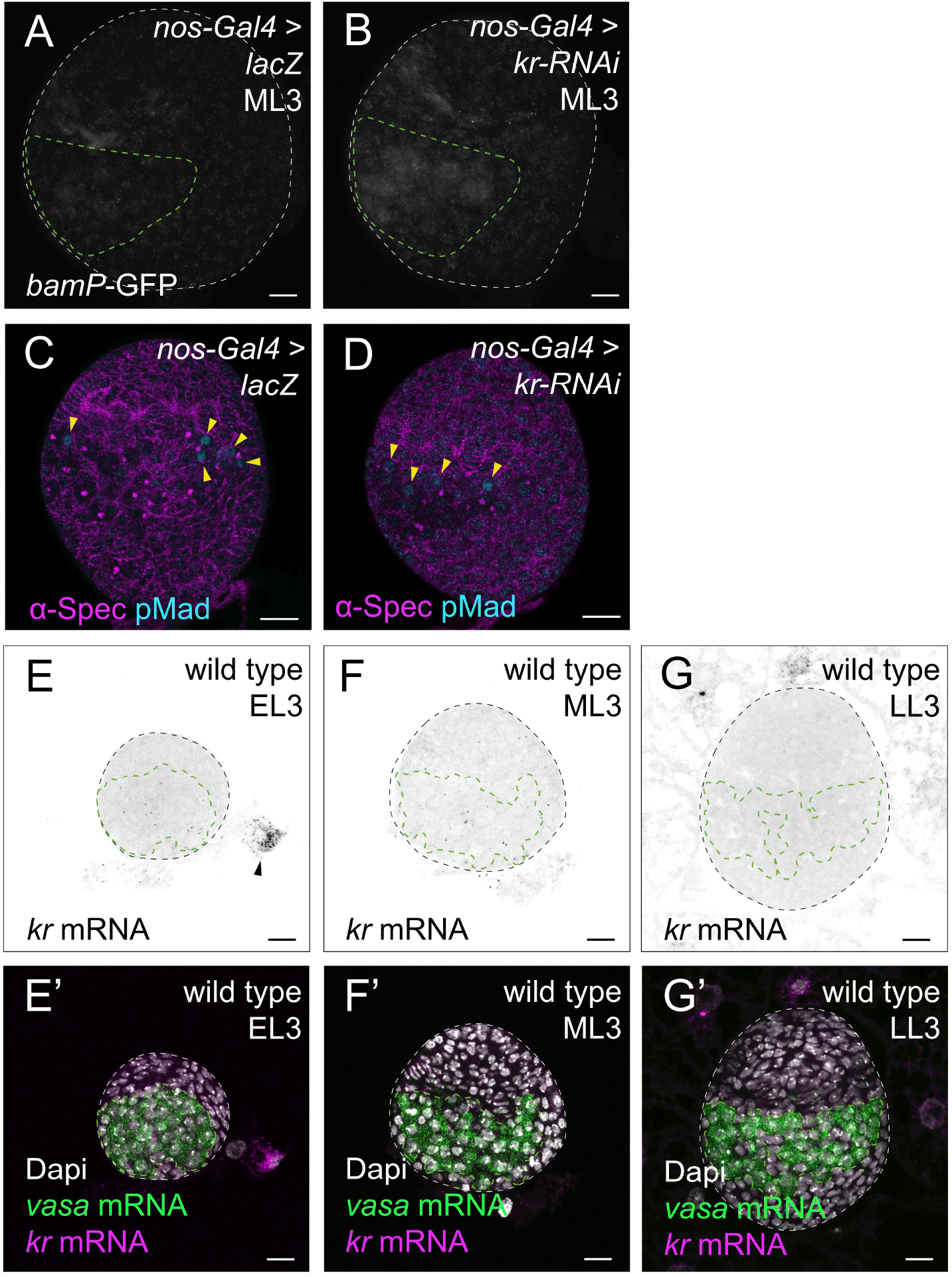
**(complementary to Figure 6) Kr represses precocious PGC differentiation** (A-B) Related to Figure 6A-D. *kr*-RNAi knockdown results in precocious PGC differentiation at ML3. PGC differentiation status is indicated by *bamP*:GFP expression; germ cell domain is outlined in green. (C-D) Related to Figure 6B. RNAi mediated knockdown of *kr* does not impact pMad expression in GSCs. (E-G) Related to Figure 6E. HCR *in situ* hybridization for *kr*-mRNA at different developmental stages; *kr*-mRNA expression in the larval fat body is indicated (asterisk). *vasa*-mRNA expression pattern indicates germ cell position (green outline); Dapi labels all nuclei. Scale bars indicate 10 μm.

## Acknowledgements

We like to thank Drs. Jordi Casanova, Michael Buszczak, Dorothea Godt, Erika Bach, Jessica Treisman, Stephen Crews and Katsuo Furukubo-Tokunaga for kindly sharing their reagents. We thank the TRiP at Harvard Medical School for providing transgenic RNAi fly stocks used in this study. We thank the Bloomington, VDRC, DGRC and FlyORF stock collections for the stocks used in this study. We acknowledge the FlyBase team for their exceptional work on the *Drosophila* database. We thank Shifra Ben-Dor at the Life Sciences Core Facilities at the Weizmann Institute of Science for *in-silico* prediction of Krüppel binding sites in the *bam* promoter. We thank Toby Lieber and Lacy Barton for critical review of the manuscript and all members of the Lehmann lab for helpful discussions, constructive comments and ideas.

## Funding

T.U.B. was supported by a European Molecular Biology Organisation long-term fellowship (ALTF47-2014). M.S. was a Howard Hughes Medical Institute (HHMI) Fellow of the Life Sciences Research Foundation. S.G. is supported by NYU Dean’s Undergraduate Research Fund Grant, L.G. was supported by Israel Cancer Research Fund (12-3073-PG) and by the Helen and Martin Kimmel Institute for Stem Cell Research at the Weizmann Institute of Science (HMKISCR). Research in the Lehmann lab is supported by HHMI, where R.L. is an investigator, and by NIH R37HD41900.

## Author Contributions

Conceptualization, R.L., L.G. and T.U.B; Methodology, R.L., L.G. and T.U.B; Investigation, T.U.B and S.G.; Software and Resources, M.S.; Writing – Original Draft, T.U.B., L.G., R.L.; Supervision, R.L. and L.G.; Funding acquisition, R.L., L.G. and T.U.B;

## Declaration of Interests

The authors declare no competing interests

## STAR Methods

### Lead Contact and Materials Availability

Information regarding requests for materials used in this paper are listed in the Key Resources Table. Further information and requests for reagents, protocols or other resources should be directed and will be fulfilled by the lead contact, Ruth Lehmann.

### Experimental Model and Subject Details

*Drosophila melanogaster* were raised on medium containing yeast, molasses and cornmeal, and kept at 25°C. A complete list of fly lines used in this study can be found in the Key Resources Table.

### Method details

#### Staging

To obtain flies of similar developmental stages, care was taken to work with under-crowded cultures. Flies were transferred into a fresh vial to lay eggs for 3h, and were then removed. Vials were left at 25° for 48h (second larval instar), 72h (early third larval instar) for 96 h (mid third instar, ML3) or 120 h (late third instar, LL3). Under these conditions the development of wildtype gonads is uniform. To further account for staging differences, all gonads were co-stained with Dapi and TF maturation status and allover ovary size was used as proxy to verify accurate staging. The terminology we use is according to Ashburner (Ashburner, 1989).

#### Dissections

Larvae were dissected as described previously (Maimon and Gilboa, 2011). In short: Larvae were rinsed in Ringer’s solution or DPBS (for *in situ* hybridization). Heads were removed with forceps. Specimens were inverted and trachea and guts were gently removed. For *in situ* and EL3 stages the fatbody and was left attached to the cuticle, for all other purposes the cuticle was removed prior to staining. Properly staged pupae were dissected following the procedures in (Park et al., 2018). For adults, females were dissected 3 days after eclosion with 1day fattening on yeast. Abdomens were removed using forceps, intestine was removed and staining was done with the ovaries partly covered by the abdominal cuticle.

#### Immunofluorescence staining

All steps were performed with gentle rotation. Specimens were fixed in Ringer’s solution, 4% Paraformaldehyde for 20min at RT, followed by a short was with 1%PBT (TritonX in PBS), and 1h wash in PBT for permeabilization. Blocking was done in 0.3% PBTB (TritonX in BSA) for at least 1h at RT. Primary antibodies were incubated at 4°C, overnight in 0.3% PBTB. Subsequently, specimens were washed twice for 30min with 0.3% PBTB, followed by 1h in 0.3% PBTB. Secondary antibodies were diluted in 0.3% PBTB and incubated for 2h at RT, followed by three washes in 0.3%PBT (TritonX in PBS). For mounting, specimens were equilibrated in Vectashield+Dapi and the fatbody was removed from the gonads with dissecting needles.

#### RNA *in situ* hybridization

All steps are done in using RNAse free reagents and supplies and with gentle rotation, except for steps at 37°C. The protocol was adapted from (Choi et al., 2018). Specimens were fixed in PBS, Tween (Tw) 0.1%, Paraformaldehyde 4% for 20 minutes at RT, washed twice with PBS, Tw 0.1% at RT, dehydrated with sequential washes with 25%, 50%, 75% and 100% methanol in PBS on ice 5 minutes each. Samples were stored at −20°C at least overnight (up to one week). Samples were rehydrated with sequential washes with 100%, 75%, 50% and 25% methanol in PBS on ice, permeated for 2 hours in PBS Tx 1% at RT, post-fixed in PBS, Tw 0.1%, Paraformaldehyde 4% for 20 minutes at RT, washed twice with PBS, Tw 0.1% for 5 minutes on ice, washed with 50% PBS, Tw 0.1%/ 50% 5xSSCT (5xSSC, Tween 0.1%) for 5 minutes on ice, washed twice with 5xSSCT for 5 minutes on ice, incubate in probe hybridization buffer for 5 minutes on ice, pre-hybridized in probe hybridization buffer for 30 minutes at 37°C, and hybridized overnight at 37°C. Probe concentrations were determined empirically, and ranged 4 −8 pmol of each probe in 1 ml, probe solution was prepared by adding probes to pre-warmed probe hybridization solution. After hybridization, specimens were washed 4 times with probe wash buffer for 15 minutes each at 37°C, and twice with 5xSSCT for 5 minutes each at RT. Specimens were equilibrated in amplification buffer for 5 minutes at RT. Hairpin solutions were prepared by heating 30 pmol of each hairpin for 90 seconds at 95°C and cooling at RT in a the dark for 30 minutes, and subsequently adding the snap-cooled hairpins to 500 μl of amplification buffer at RT. Specimens were incubated in hairpin solution overnight at RT, and washed multiple times with 5xSSCT – twice for 5 minutes, twice for 30 minutes and once for 5 minutes. Dapi was added in the first 30-minute wash. Specimens were either equilibrated in Vectashield overnight at 4°C and mounted in Vectashield, or subsequent immunofluorescence staining was carried out (see above).

#### Detection of mRNA foci

After the HCR *in-situ* protocol (see above), gonads were scanned on the confocal microscope, a volume of 5 μm in 1 μm z-plane step was acquired. Stacks were maximum projected in Fiji software. Area of SwCs (*sim* signal) or PGCs (*vasa* signal) was outlined and area around this ROI removed (Fiji edit>clear outside). Nuclei contained in the area were counted manually (Dapi signal) to determine SwC/PGC numbers. mRNA abundance was determined using the freely available spot-detection algorithm (Airlocalize) as described in (Trcek et al., 2017), which was developed in the MATLAB programming language (MathWorks) by (Lionnet et al., 2011). Briefly: batch mode analysis of 2d images was performed. To increase spot detection accuracy initial settings for the analysis were determined on a smaller ROI in an image file; a generated image of such detected spots was overlaid with the original image file to confirm accuracy. An intensity histogram of identified mRNA spots was plotted, and a mean intensity value determined by Gaussian distribution and by verification of high and low values to actual mRNA spots in the image file. Spots around the mean intensity (1): > 0.5 = 1 < 1.5 were counted as single mRNA foci; < 0.5 as background and > 1.5 stepwise as 2, 3 foci etc. Notably, mRNA spots detected *via* the software likely present single RNA molecules, however since HCR amplifies the signal from used probes we refer to these spots as mRNA foci. Since we assume that HCR amplification is even across detected mRNAs we assign higher than mean pixel intensity values to more than one mRNA foci. The number of mRNA foci obtained for a single ovary was divided by the number of cells (SwCs or PGCs) counted for this particular ovary to obtain the plotted mean per cell value.

#### Imaging and image analysis

Images were acquired using Zeiss LSM 800 and 780 confocal microscopes using 40x oil NA 1.3 objectives. Confocal images shown are maximum projections of 3z-planes 1μm apart.

#### Lineage tracing

Lineage specific expression of *sim*-Gal4 and *cv-2*-Gal4 was validated by expression of a G-TRACE cassette. Positively marked cells were followed from L2 to late pupal stages. To lineage trace SwCs marked at LL3 into adult stages, temperature shifts utilizing the Gal80ts system were performed. Larvae were raised in the restrictive temperature (25°) till 120h-144h ael and subsequently shifted to the permissive temperature (18°). Adults were dissected 1d after eclosion.

#### Swarm cell motility index

Gonads were dissected and IHC performed at 72h, 96h and 102h. Subsequently, SwCs positively marked by *sim-G4>Eb1:GFP* were analyzed. For each SwC, ‘motile morphology’ was determined, where a motile cell is defined as having both, cellular protrusions generated towards the gonad posterior and elongated nuclear morphology. Both parameters were scored visually, with Dapi marking nuclei and Eb1:GFP labeling protrusions. 8% of analyzed SwCs displayed only one migratory parameter and were scored as non-motile. These might be stationary cells with protrusions generated in random directions or migratory cells in the process of reorienting their motile machinery.

#### qPCR analysis

Larva dissections were carried out as described previously (Gancz and Gilboa, 2017). Gonads were frozen in liquid nitrogen and stored at −80°C for one day. For best RNA yield, 15-20 gonads of LL3 and 25 gonads of ML3 were used. RNA extraction was done using RNeasy micro Kit from Qiagen following manufacturers protocol; followed by cDNA synthesis with Supercript IV from Invitrogen. mRNA levels of *rps17* served as a control. To further test whether temporal control via ecdysone was limited to *tsl* mRNA expression, qPCRs for *trunk*- and *torso* mRNA levels showed no change from ML3 to LL3 (Figure S2H). qPCR analysis was performed on a Roche LightCycler 480 II system.

#### Differentiation assay – *bam*P:GFP levels

Ovaries for wildtype and experiment were dissected, stained and imaged at the same time with the same settings throughout. For confocal imaging, a volume encompassing all germ cells was acquired. Subsequent analysis was performed in Fiji. The ten brightest germ cells per gonad were determined visually; ROI outlined in Fiji and mean pixel intensity values measured. The plotted relative intensity values represent the ratio of measured values and the mean wildtype value of the same experiment. For each experiment up to 10 gonads were imaged and each experimental condition was repeated at least three times.

### Quantification and statistical analysis

All experiments were performed with at least three biological repeats. For all experiments, over 25 ovaries were examined. Statistical significance was determined using Mann-Whitney *U*-test. Error bars represent standard deviation.

### Data and code availability

The single cell RNA sequencing data set used in this study has been published and all associated data are available in (Slaidina et al., 2020).

## Notes

### Competing Interest Statement

The authors have declared no competing interest.

